# Different developmental trajectories of critical dynamics in prefrontal cortex–amygdala circuitry

**DOI:** 10.64898/2026.02.13.705704

**Authors:** Kharybina Zoia, Palva Matias, Palva Satu, Lauri Sari, Hartung Henrike, Taira Tomi

## Abstract

Neuronal avalanches, a hallmark of self-organized criticality, represent cascading activity among interconnected neurons. Despite their hypothesized role in neuronal maturation, developmental patterns of neuronal avalanches across interconnected brain regions *in vivo* remain unelucidated. We applied avalanche analysis to investigate the emergent features of functional connectivity in the developing medial prefrontal cortex–basolateral amygdala (mPFC–BLA) circuitry, using simultaneous electrophysiological recordings from the BLA and mPFC in juvenile and young adult anaesthetized rats. The avalanches were mainly confined to either mPFC or BLA, a small number spanned both regions. Prefrontal avalanches exhibited scale-free behavior with branching ratio close to one and decreasing over development; amygdaloid avalanche size distributions displayed a sharp peak at the maximum size, more pronounced in adulthood, which together with branching ratios over one suggests a deviation from criticality toward a supercritical state. Different dynamical regimes in the BLA and mPFC fit distinct functional demands of these regions.

## Introduction

The cortico-limbic connections that support emotional behavior develop over a prolonged period early in life. This development unfolds over an extended period, marked by asynchronous maturation of the two systems, and is essential for achieving adult-level emotional and cognitive regulation (Yang and Tseng, 2022).

During this period, corticolimbic synaptic connections are gradually refined, a process essential for the development of mature emotional regulation, decision-making, memory, and social behavior. Adverse experiences in early life can alter the trajectory of corticolimbic development, leading to abnormal connectivity, such as hyperinnervation of certain limbic-to-cortical circuits which may contribute to heightened susceptibility to anxiety, depression, and other emotional or behavioral problems later on (Braun and Bock, 2011; Haikonen *et al*., 2022; Donati, Vedele and Hartung, 2024).

It is known that a critical factor for a given set of neurons to become functionally connected is the temporal coincidence of their activity (Katz, 1993; Turrigiano and Nelson, 2004; Huupponen *et al*., 2013; Turrigiano, 2017). Consequently, the width of the temporal integration window defines the minimum rate at which populations can be dissociated, a narrower integration window allowing for a larger number of independent coincidences each time. The power-law distribution of neuronal avalanches indicates that the brain is capable of processing neuronal signaling across a wide range of spatial (millimeter-centimeter) and temporal (millisecond-hours) scales (Shriki *et al*., 2013; Tomen, Rotermund and Ernst, 2014). By this virtue, neuronal avalanches can be instrumental in both the Hebbian and homeostatic plasticity involved in strengthening and weakening synaptic connections during brain development. Moreover, Ma et al. (Ma *et al*., 2019) showed that after perturbed activity and the resulting deviation from critical dynamics in the visual cortex, homeostatic plasticity in inhibitory synapses re-establishes critical dynamics in excitatory networks. Thus, neuronal avalanches may both guide and reflect the developmental status of the neuronal network.

Neuronal avalanches have been proposed to be involved in emergence of efficient network topologies (Shew *et al*., 2011), synaptic pruning and refinement (Michiels Van Kessenich, De Arcangelis and Herrmann, 2016; Yada *et al*., 2017), cortical specialization (Gireesh and Plenz, 2008) and cognitive development (Chialvo, 2010). Given the protracted development and highly plastic nature of developing corticolimbic circuitry, neuronal avalanches could provide an effective endogenous drive for corticolimbic synaptic refinement. An important notion here is that the brain exhibits a hierarchical, modular organization in which groups of neurons form functional clusters. It has been suggested that networks with modular organization preserve criticality more robustly than those that are fully interconnected (Rusch, Kinouchi and Roque, 2025). Two important consequences follow from this. First, such an organization allows activity to remain contained within a module rather than spreading uncontrollably across the entire network. Second, it means that avalanche dynamics and synapse formation can differ markedly even between closely interconnected brain regions.

The intrinsic activity patterns that are linked to the development of limbic circuits remain poorly understood. Given the known differences between the mPFC and BLA in both functional architecture and physiological roles (Sah *et al*., 2003; Dalley, Cardinal and Robbins, 2004; Petrides, 2005; Phelps and LeDoux, 2005; Janak and Tye, 2015; Anastasiades and Carter, 2021; Wassum, 2022), we were intrigued to explore the critical dynamics in these brain regions during development. Here we have examined how the dynamical states of the mPFC–BLA circuitry change over normal development in a rat model using neuronal avalanche analysis (Beggs and Plenz, 2003; Petermann *et al*., 2009).

## Materials and Methods

### Animals

All experiments were approved by the National Animal Experiment Board in Finland, ELLA, and adhere to the University of Helsinki Animal Welfare Guidelines.

Experiments were done on Han-Wistar rats, both males and females, in two age groups: juvenile (p14-p15) and young adult (p50-p60). Pups were weaned on p21 and then group housed (2-3 animals of the same sex in a cage) under standard housing conditions (temperature and humidity-controlled environment, ad libitum access to food and water, 12/12h light/dark cycle). The Estrous cycle stage was not controlled.

Animals from the same liters were randomly divided into juvenile and young adult groups. In total, animals from 19 liters were used for experiments. We recorded from 48 young adult animals from which 6 animals were excluded (one animal died during recording, two recordings had corrupted signal, and three recordings had too large noise caused by a heating pad), and 36 juvenile animals from which two died during recording and one was excluded because of the corrupted signal. Depending on the analysis, individual recordings were excluded due to the insufficient number of channels in the region of interest or low signal to noise ratio (MUA).

### In vivo electrophysiology

#### Surgical procedure and recordings

We performed extracellular recordings in the basolateral amygdala (BLA) and prelimbic division (PL) of the medial prefrontal cortex (mPFC) of urethane-anaesthetized rats. After anaesthesia induction with 4% isoflurane, urethane (10% urethane diluted in NaCl, 13 µl/g, i.p. injection) was administered at least 40 – 50 minutes before recordings. Surgery was performed under continuously administered 1 – 2% isoflurane for surgically deep anaesthesia. 3 burr holes were drilled: 2 above regions of interest (BLA: base of the rhinal fissure, PFC: 0.5 – 1.5 mm and 1.5 – 2.5mm anterior to bregma for juvenile and young adult animals, respectively, 0.3 mm from the midline) and 1 above cerebellum for reference electrode. Then, the animal was placed into the stereotaxic frame, and the head was fixed using ear bars for young adult animals or bars fixed with dental cement on the nasal and occipital bones for juvenile animals.

For recordings, we used 32-channel four-shank (4×8) or 64-channel eight-shank electrodes (8×8) silicon multielectrode probes (NeuroNexus Technologies) with 200 µm distance between shanks and 200 µm pitch between the recording sites. These distances between channels allow to avoid spurious correlations in the nearby channels (Neto, Spitzner and Priesemann, 2022). A four-shank probe was inserted into the mPFC perpendicularly to the skull surface and parallel to the midline to have all the shanks in the same layer, 3.5 mm depth. Into the BLA, a probe was inserted at 40° from the vertical plane. For most young adult animals, we used an eight-shank probe, for the rest of the young adult animals and juvenile animals we used a four-shank probe. The depth was 5 – 5.5 mm and 4.3 – 4.7 mm for young adult and juvenile animals, respectively. For postmortem electrode track reconstruction, the electrodes were labeled with DiI (1,1’-dioctadecyl-3,3,3’,3’-tetramethyl indocarbocyanine, Invitrogen). A common for both head stages silver wire reference electrode was inserted into the cerebellum. During surgery and recordings, the body temperature was maintained at 37°C with a heating pad. If needed, a urethane top-up of 1/3 of the original dose was administered.

After probe insertion, animals recovered for 15 min. Then simultaneous recordings of local field potential (LFP) and multi-unit activity (MUA) were acquired with 20 or 32 kHz sampling rate for 45 minutes. For recordings, we used Digital Lynx 4SX, Neuralynx and Open Ephys acquisition board, Open Ephys. Each animal was recorded only once and was sacrificed after the recording.

#### Histology

After the recording, animals were perfused with 4% paraformaldehyde (PFA), brains were extracted and stored for at least 3 days in PFA. Then the brains were dissected (100-µm thick slices, coronal plane), sections in the regions of interest were stained using Nissl staining procedure, imaged and used for electrode reconstruction.

### Data analysis

For data analysis, we used only electrodes confined in the BLA and PL confirmed by postmortem histological examination. Experiments with channels in deep and superficial layers of the mPFC were pulled together. Data was imported and analyzed in MATLAB (R2020b, Mathworks). Signal traces were low-pass filtered (third order Butterworth filter, < 1000 Hz), downsampled to 2 kHz and notch-filtered at 50 Hz (IIR notch-filter, q-factor = 35) with MATLAB *filtfilt* function to avoid phase distortions.

#### Avalanche detection

For avalanche detection, we picked 16 channels in a region of interest. Recordings with a smaller total number of channels in a region were not used in the analysis. Avalanches were calculated separately for the mPFC and the BLA. Number of animals: juvenile group mPFC 15 males and 15 females, BLA 11 males and 17 females; young adult group mPFC 17 males and 15 females, BLA 13 males and 15 females.

To obtain LFP, time courses signals were normalized (demeaned and divided by signal standard deviation) and band-pass filtered (third order Butterworth filter, MATLAB *filtfilt* function, 1 – 100 Hz and 2 – 100 Hz for BLA and mPFC traces, respectively). We used higher low cut-off frequency in the case of the mPFC to avoid an effect of anaesthesia-induced high amplitude slow-rhythm prominent in this region.

LFP time courses from 16 channels were converted into spatio-temporal clusters of negative LFP deflections (nLFPs), or neuronal avalanches. nLFPs were defined as maximum negative LFP deflections upon crossing a threshold. Thresholds for nLFP detection were calculated based on the standard deviation *SD* of an individual trace. All calculations were done for threshold values starting from 1*sd* up to 4*SD* with the step of 0.2*SD*. Thresholds exceeding 4*SD* resulted in a low number of nLFPs, insufficient to form spatio-temporal clusters.

nLFP rasters were divided into time bins. Calculations were performed for different values of the bin length *Δt* in the range of 1 – 30 ms with the step of 0.2 ms. Subsequent bins with nLFPs, preceded and followed by at least one empty bin, formed one avalanche.

For each avalanche, we calculated its size *s* as a number of nLFPs comprising it and branching ratio *σ*, which is the ratio of nLFP counts in the second and first bins of the avalanche. Then we calculated avalanche size distributions.

We fitted size distributions (from 1 to 16 channels) with truncated power-law function *P(s)∼e^-αs^s^τ^* using the least-squares method and estimated the goodness of fit by calculating R squared.

For mean avalanche profiles we calculated the number of nLFPs in every bin of the avalanche and averaged it over all avalanches with the same duration. Avalanches with durations less than 4 and larger than 9 bins were excluded from analysis since in the short avalanches the number of bins was too low to estimate the shape of the avalanche, whereas long avalanches were infrequent and had large variation in the shape.

For region comparison and for avalanche spread analysis, we used only a fraction of experiments where we had at least 16 channels in both regions simultaneously. Number of animals: juvenile group 11 males and 14 females; young adult group 11 males and 11 females.

#### Power spectrum and functional connectivity

For spectral and connectivity analysis, we performed a continuous wavelet transform of the voltage time series (complex Morlet wavelet, length 5 cycles, 50 center frequencies logarithmically distributed in the band 0.1 – 100 Hz). A square of the absolute value of wavelet transformed signal was converted to decibel to obtain spectral power.

To assess functional connectivity between two channels we calculated phase locking value (PLV) as

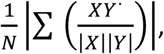

where *X* and *Y* are wavelet transformed voltage signals, *Y\** - complex conjugate of *Y*, *N* – length of the signal.

For spectral and in-region functional connectivity analysis, we used the same 16 channels as in avalanche calculations and presented results as average values for all channels (power) or pairs of channels (PLV). For functional connectivity between the regions, we used only recordings with channels in both regions and selected one channel per region based on histology and signal to noise ratio. Group sizes in the latter case: juvenile group 14 males and 14 females; young adult group 21 males and 15 females.

To assess confidence limits for PLV we used Monte Carlo simulation. We calculated PLV between original voltage signals in the mPFC and time shuffled (3 s windows) signal from the BLA. This procedure was repeated 100 times, and the threshold was determined as 95^th^ percentile of the resulting distribution.

#### Detrended fluctuation analysis

For DFA analysis, we selected 5 nearby channels based on histology and signal-to-noise ratio. Results were averaged over these channels. Number of animals: juvenile group mPFC 15 males and 15 females, BLA 15 males and 18 females; young adult group mPFC 21 males and 19 females, BLA 18 males and 18 females.

Voltage signal underwent wavelet transform first (complex Morlet wavelet, length 5 cycles, 20 center frequencies logarithmically distributed in the band 1 – 100 Hz), followed by Hilbert transform. Then we calculated a cumulative sum of the transformed signal to get a signal profile.

A fluctuation function of the signal profile was calculated in the following way. Time series were divided into the windows with 80% overlap. For each window we calculated standard deviation of the detrended signal and then calculated mean standard deviation for all the windows. This procedure was repeated on the set of windows with sizes logarithmically distributed from 1s to 500s (up to 20% of the signal length).

A fluctuation function, in other words a mean standard deviation of the detrended signal as a function of window size, follows power law. To get DFA exponents we calculated a slope of fluctuation function in double-logarithmic scale using least-squares fit. To avoid effects of filter-caused distortions in case of small window sizes and low number of windows for larger sizes, we used an interval from 5s to 300s for linear fit.

#### Multi-unit activity

To detect multi-unit activity, we picked one channel in the region of interest with the most prominent spiking. Number of animals: juvenile group mPFC 12 males and 12 females, BLA 12 males and 15 females; young adult group mPFC 17 males and 15 females, BLA 12 males and 15 females.

Signal was filtered (Butterworth high-pass filter with cut-off at 300 Hz). For spike detection threshold we used an estimate of the standard deviation of the background noise

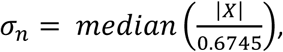

where *X* is a filtered signal (see (Quiroga, Nadasdy and Ben-Shaul, 2004)). The threshold was set as *3.5σ_n_*.

#### Statistical analysis

Developmental changes in the power of oscillations, PLV and DFA were assessed with separate statistical tests for each frequency (Wilcoxon ranksum test). To compare branching ratios, separate statistical tests were run for each pair of parameters (threshold and bin size). We used Wilcoxon ranksum test to assess differences between age groups and Wilcoxon sign-rank test for paired observations in case of regional differences. False discovery rate was controlled with the Benjamini-Hochberg procedure, accepted q-values less 0.05.

To assess the effect size, we calculated probabilistic index

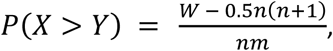

where *W* is the Wilcoxon statistics, *m* and *n* – number of observations in the groups *X* and *Y*. Probabilistic index was calculated for all points in the parameter space and averaged over all points belonging to the same clusters of points with significant differences on the color-coded heat map.

Two-way repeated measures ANOVA with Holm-Sidak post-hoc all pairwise multiple comparisons was used to assess developmental changes in avalanche localization.

Developmental changes in MUA activity were assessed either with Welch’s t-test when normality assumption was passed or with Wilcoxon rank sum test otherwise.

## Results

### 1. Oscillation power and functional connectivity in the mPFC and BLA increase over development

To investigate developmental changes in the neuronal dynamics of the mPFC-BLA circuit we performed simultaneous multi-electrode *in vivo* local-field potential (LFP) and multi-unit activity (MUA) recordings (45 min) from the BLA and the prelimbic part of the medial PFC (Figure 1A, 1F) in lightly urethane-anesthetized male and female rats during either juvenile (postnatal day (p) 14 – p15, N = 36)) or young adult (p50 – p60, N = 48) development. In juvenile animals, activity was already continuous; the predominant state throughout the recording was slow-wave sleep where high-amplitude slow-wave activity (0.2 – 0.5 Hz) was superimposed by faster rhythms. In adult animals, one third of the recordings showed the typical brain-state changes between high-amplitude slow-wave sleep (∼1 Hz) and faster activity dominated by delta/low-theta activity (2 – 4 Hz) as previously described in adult rodents (Clement *et al*., 2008; Silver, Ward-Flanagan and Dickson, 2021; Ward-Flanagan *et al*., 2022), however, brain state shifts pattern lacked regularity. Typically, one of these states was predominant and the other state only contributed to 10 – 20% of the recording. In the other two thirds of the recordings only one state was present. In both age groups, anesthesia-induced high-amplitude slow-wave activity was more prominent in the mPFC than in the BLA.

**Figure 1.**
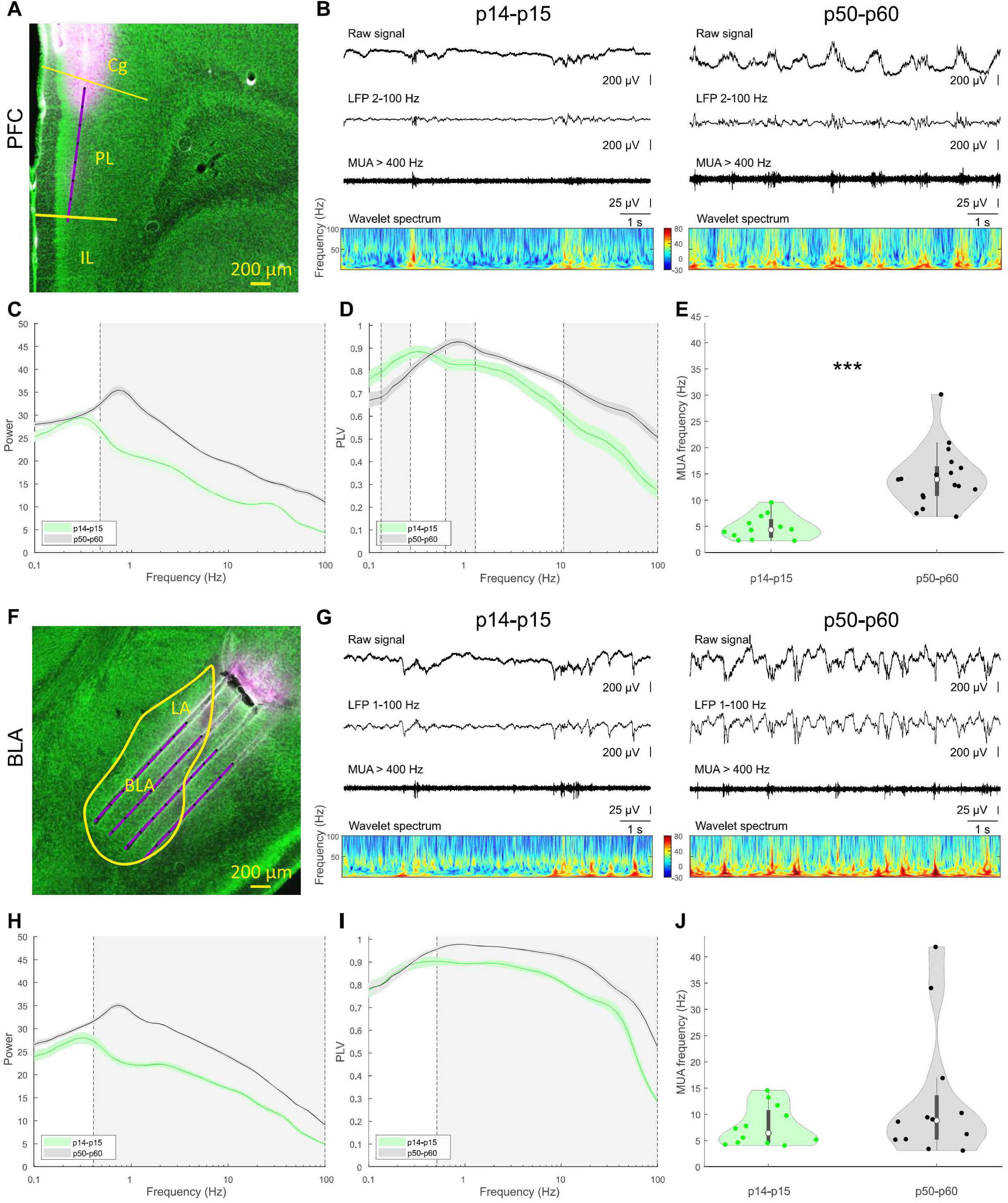
LFP signal in the mPFC and the BLA in male rats. (A, F) Digital reconstruction of the electrode positions in Nissl-stained 100-µm coronal sections from the mPFC (A) and the BLA (F). In the mPFC, the red line indicates the recording position of a single electrode shank of the multi-shank probe positioned parallel to the midline. In the BLA, red lines indicate positions of all four electrode shanks covering all subregions of the BLA. Black dots represent individual recording sites. For the avalanche analysis, 16 channels within the PL subregion of the mPFC were chosen as well as 16 channels within the boundaries of the BLA. (B, G) Extracellular LFP and MUA recording of neuronal activity in the mPFC (B) and BLA (G) from a juvenile p14 pup (left panel) and young adult p50 rat (right panel). From top to bottom: raw trace, band-pass filtered signal 2 – 100 Hz or 1 – 100 Hz for the mPFC and the BLA, respectively, corresponding MUA, obtained with 400 Hz high-pass filtering, and wavelet spectrum for the same time period. Signal traces are in µV. (C, H) Mean power spectra calculated for 16 channels in the mPFC (C) and the BLA (H) from juvenile (green, N = 15, 11 for the mPFC and the BLA, respectively) and young adult (black, N = 17, 13 for the mPFC and the BLA, respectively) rats. Shaded area shows SEM. Grey rectangles represent significant differences (p < 0.05, Wilcoxon rank-sum test with FDR correction, q < 0.05). (D, I) Same as (C, H) for phase locking value (PLV). (E, J) Instantaneous MUA firing rate (Hz) calculated for a single channel in the mPFC (E) and the BLA (J) of juvenile (green, N = 12 for both mPFC and BLA) and young adult (black, N = 17, 12 for the mPFC and the BLA, respectively) rats. Firing rates increase with age in the mPFC (juveniles: median (IQR) = 4.35 (3.44), adults: median (IQR) = 13.95 (5.59), p = 2.32E-06, Welch’s t-test), but remain similar across development in the BLA (juveniles: median (IQR) = 6.43 (6.14), adults: median (IQR) = 8.82 (8.35), p = 0.47, Wilcoxon rank sum test).

For each region we calculated a wavelet power spectrum as an average for 16 adjacent channels. In both mPFC and BLA, power significantly increased with age in a broad band (Figure 1C, 1H, S1A, S1D, mPFC: 0.5 – 100 Hz, n = 15, 17 for juvenile and adult males, respectively, BLA: 0.4 – 100 Hz, n = 11, 13, for juvenile and adult males, respectively, p < 0.05, Wilcoxon rank sum test with false discovery rate (FDR) correction). As expected under urethane anesthesia (Clement *et al*., 2008), slow-wave rhythmicity prevailed in the recordings, with peak frequencies significantly shifting towards higher values with age in both brain regions (mPFC, juvenile males: 0.29 ± 0.01 Hz, n = 15, young adult males: 0.69 ± 0.04 Hz, n = 17, p = 9.74E-06; BLA, juvenile males: 0.32 ± 0.01 Hz, n = 11, young adult males: 0.72 ± 0.06 Hz, n = 13, p = 4.15E-4, Wilcoxon rank sum test).

To assess pairwise functional connectivity between the same 16 channels within either the mPFC or the BLA, we computed phase locking value (PLV) and averaged it over all pairs of the channels. Due to the small distance between adjacent electrodes (200 µm), the signal in all channels shared high similarity resulting in high PLV values close to 1. In males, PLV in the mPFC (Figure 1D) significantly decreased with age for ultra slow frequencies (0.13 – 0.27 Hz) and increased for higher frequencies in broad bands (0.6 – 1.3 Hz and 10 – 100 Hz, n = 15, 17, for juvenile and young adult males, respectively, p < 0.05, Wilcoxon rank sum test with FDR correction). In the BLA (Figure 1I) we saw developmental increase in PLV for most of the analyzed frequencies (0.84 – 100 Hz, n = 11, 13, for juvenile and young adult males, respectively, p < 0.05 Wilcoxon rank sum test with FDR correction). In females, we observed similar, though less pronounced, developmental changes in the functional connectivity as in males (Figure S1B, S1E).

Next, we analyzed multi-unit activity (MUA). While the mPFC typically showed pronounced spiking activity in the majority of recorded channels, BLA spiking was rather sparse with only a few channels showing any MUA at all. For this reason, we limited the MUA analysis to a single channel with the best signal-to-noise ratio from the mPFC and/or BLA from each pup and discarded experiments with insufficient amplitude of spikes. MUA frequency increased with age in the mPFC (Figure 1E, S1C, juvenile males: median (IQR) = 4.35 (3.44), n = 12, young adult males: median (IQR) = 13.95 (5.59), n = 17, p = 2.32E-06, Welch’s t-test), whereas in the BLA no developmental difference was observed (Figure 1J, S1F juvenile males: median (IQR) = 6.43 (6.14), n = 12, young adult males: median (IQR) = 8.82 (8.35), n = 12, p = 0.47, Wilcoxon rank-sum test).

In summary, during ongoing development of mPFC and BLA power of oscillations as well as functional connectivity increased within these regions. In addition, multi-unit activity increased with age in mPFC but not in BLA. These effects were similar in both males and females though less pronounced in females.

### 2. Dynamical state in the mPFC changes over development

The developmental changes in network activity of the mPFC and the BLA may reflect changes in the underlying dynamical states of these regions. We investigated brain dynamics by characterizing neuronal avalanches and long-range temporal correlations in these regions.

#### 2.1. Prefrontal neuronal avalanches display scale-free behavior with branching ratio decreasing over development

To assess brain dynamical state, we performed avalanche analysis. For this, we calculated negative deflections of local field potential (nLFPs) for adjacent 16 recording channels (Figure 2A). In both mPFC and BLA, most of the nLFPs did not occur individually or simultaneously in all channels but rather formed spatiotemporal clusters, or neuronal avalanches (Figure 2B). Avalanches were defined based on the temporal order of nLFPs irrespective of the spatial position of the electrodes nLFPs occurred on. According to such definition, an avalanche comprised of all successive time bins containing at least one nLFP, was preceded and followed by at least one empty bin. As expected, avalanche characteristics depended on the bin size and threshold for nLFP detection. For this reason, we explored a wide range of parameters: bin size 1 – 30 ms and threshold 1 – 4 SD. Higher values of threshold resulted in insufficient number of nLFPs.

**Figure 2.**
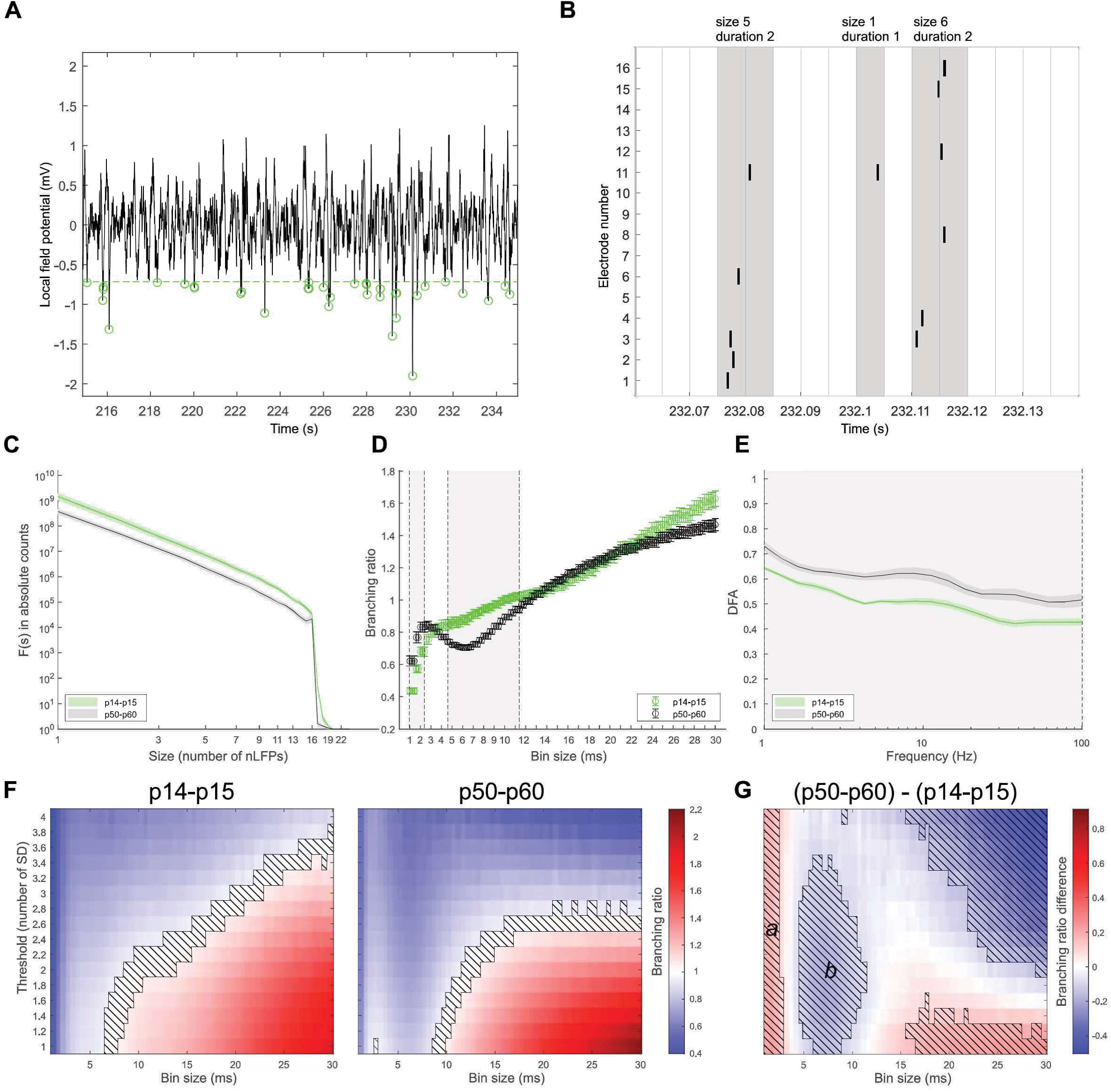
Neuronal avalanches and LRTCs across development in the mPFC in male rats. (A) An example of a band-pass filtered (2-100 Hz) LFP recording (20 s) from a p14 pup illustrating the detection of nLFPs (green circles) by applying a 2SD threshold (green dashed horizontal line). (B) Occurrence of nLFPs from individual electrodes over time within 5 ms bins. Note that in this example nLFPs occur in three avalanches (consecutive time bins with nLFPs (grey bins) separated by empty bins (white bins)). (C) Avalanche size distribution calculated with 2SD threshold and 2 ms time bin and averaged across animals (N = 15, 17 for juvenile and adult animals, respectively). In log-log scale, distribution represents straight line with a cut-off at 16 channels and is described by truncated power law. (D) Mean branching ratios calculated with 2SD threshold for multiple time bins and averaged across animals (N = 15, 17 for juvenile and adult animals, respectively). (E) Mean DFA exponents calculated for five nearby channels and averaged across animals (N = 15, 21 for juvenile and adult animals, respectively). In (C-E) results for juvenile and adult animals are shown in green and black colors, respectively. Shaded area and error bars show SEM. Grey rectangles correspond to significant differences (p < 0.05, Wilcoxon rank sum test with FDR correction, q < 0.05). (F) Mean branching ratios calculated with multiple thresholds and time bins and averaged across animals for juvenile (left panel) pups and young adult (right panel) rats. Hatched area corresponds to the branching ratios close to 1 (0.95 – 1.05). (G) Difference in the mean branching ratio between young adult and juvenile animals. Hatched areas correspond to the statistically significant differences (p < 0.05, Wilcoxon rank sum test with FDR correction, q < 0.05). Significant cluster ***a***: size s = 138, mean probabilistic index <*P(adult > juvenile)*> = 0.83. Significant cluster ***b***: size s = 331, mean probabilistic index *<P(adult < juvenile)>* = 0.83.

Initially, we calculated neuronal avalanches separately for different brain states (n= 3 per group). Though avalanche characteristics differed between the states, results obtained for the whole recording without distinguishing between brain states were similar to the results obtained only for the predominant state. For this reason, we ran all calculations on the whole recording without dividing it into separate states.

For each avalanche, we estimated its size *s*, or a number of nLFPs in the avalanche, and then calculated size distribution for all avalanches in the recording. In agreement with previous research on cortical avalanches (Beggs and Plenz, 2003; Gireesh and Plenz, 2008; Petermann *et al*., 2009), in the mPFC for most of the parameters avalanche size distributions were finely described by truncated power law. We estimated the goodness of fit for truncated power law calculating R-squared, and for both age groups R-squared averaged over all animals was higher than 0.9 (Figure S2) except for a small set of parameters in case of adult animals (bin sizes > 10 ms in combination with a small threshold < 1.6 SD). In double-logarithmic scale avalanche size distributions were represented by a straight line with a cut-off at the size of 16 channels (Figure 2C, S2A), the maximum number of channels used in the avalanche analysis, meaning that one recording site usually participated in the avalanche only ones. The cut-off was more prominent for smaller bin sizes; for larger bin sizes distributions were characterized by heavier tails, and a small bump appeared at the size of 16 channels though this did not affect power-law fit drastically.

For each avalanche, we calculated a branching ratio *σ*, which reflects a number of electrodes activated following an activation of a single electrode. For critical processes, on average this value should be close to one. In a supercritical state, a number of activated channels will grow over time and *σ* > 1, whereas a subcritical state with activity diminishing over time is characterized by *σ* < 1.

In the mPFC, there was a clear dependence of the branching ratio on the bin size (Figure 2D, 2F, S3B, S3D). In agreement with previous studies (Beggs and Plenz, 2003; Priesemann *et al*., 2013; Zhigalov *et al*., 2015), in both age groups the branching ratio increased with bin sizes. Interestingly, whereas in juvenile animals the branching ratio increased monotonically, unexpectedly in young adults there was a small drop in the branching ratio between 2 and 10 ms. In juveniles, for almost all threshold values there was a certain bin size producing branching ratio close to one (Figure 2F, S3D, shaded band). In young adults, this band shifted towards higher values of the bin size and was restricted by the thresholds < 3 SD.

In males, a comparison of two age groups revealed four separate clusters of parameters (Figure 2G) with significant changes in the branching ratio (p < 0.05, n = 15, 17, for juvenile and adult animals, respectively, Wilcoxon rank sum test with FDR correction). However, only clusters ***a*** and ***b*** spanned almost all threshold values and we would focus on them. To assess an effect size, we calculated probabilistic index *P(adult>juvenile)*, as described in (Acion *et al*., 2006), which shows the probability of the branching ratio in adult animals being higher than the branching ratio in juvenile animals, or vice versa *P(adult<juvenile)*, and averaged it over all points in the parameter space belonging to the cluster. For very small values of the bin size (< 3 ms, cluster ***a***) branching ratio increased with age with mean value <*P(adult>juvenile)>* being 0.83. For the bin sizes between 5 and 10 ms (cluster ***b***) we saw an opposite effect, branching ratio decreased with age, <*P(adult>juvenile)>* = 0.83. In females the trend was the same (Figure S3E), however an effect was significant only for the cluster **a** (increase in the branching ratio, small bin sizes < 3 ms) whereas a decrease in the branching ratio for the bin sizes between 5 and 10 ms did not reach level of significance.

Theory of self-organized criticality predicts that avalanche mean temporal profiles should be uniform across different time scales. In other words, when brought to a universal scale, the shapes of the avalanches should collapse (Friedman *et al*., 2012; Jenkinson and Goutsias, 2014). We checked this property on randomly selected animals from both age groups and calculated mean temporal profiles for the avalanches with the threshold of 2 SD and bin sizes from 1 to 30 ms with the step of 1 ms. We took into consideration only avalanches with durations from 4 to 9 bins since avalanches with durations 10 bins and more were rare and had high variation in the shape. Shape collapse was observed in the adult group in a range of 15 – 25 ms (Figure S4C), whereas in the juvenile group rescaled profiles aligned to a lower extent (Figure S4A).

#### 2.2. Long-range temporal correlations in the mPFC emerge only in young adulthood

To further investigate brain dynamics, we assessed long-range temporal correlations (LRTCs) by performing detrended fluctuation analyses (DFA)(Hardstone *et al*., 2012). Random processes with no memory of the previous events are described by DFA exponents close to 0.5, whereas higher DFA exponents (0.5 < *a* < 1) reflect positive LRTCs in the trace. Critical systems are characterized by maximized information storage (Haldeman and Beggs, 2005; Beggs, 2022), hence in the critical state LRTCs reflected in DFA exponents are also maximized. For DFA analysis, we used 5 adjacent channels from each region and averaged results over them. In juvenile group, except for very slow oscillations (< 2 Hz), DFA exponents in the mPFC were close to 0.5, showing an absence of LRTCs in the trace (Figure 2E, S3C). Positive LRTCs in the mPFC emerged only in young adults, where DFA exponents were significantly higher compared to juvenile animals for the whole frequency range in males (p < 0.05, n = 14, 21, for juvenile and adult animals, respectively, Wilcoxon rank sum test with FDR correction). The same developmental increase in DFA was seen in females, though it was not significant for the frequencies near 20 Hz (Figure S3C).

Summing up, in both age groups prefrontal nLFPs were clustered into neuronal avalanches with power-law size distribution as expected of the systems near the critical state. Mean branching ratio for the avalanches showed developmental increase for very small bin sizes and decrease for the bin sizes between 5 and 10 ms. The decrease was significant only in males. Prefrontal voltage traces in juvenile animals lacked LRTCS which emerged only in early adulthood where DFA exponents significantly increased compared to juvenile animals.

### 3. Developmental changes in the dynamical state in the BLA follow different trajectory from the mPFC

#### 3.1. Neuronal avalanches in the amygdala do not follow power law with branching ratio over one at both developmental stages

In contrast to the mPFC, avalanche size distributions in the BLA were characterized by a sharp peak corresponding to the maximum number of electrodes used in the analysis (Figure 3A, S5A). This means that avalanches tended to span the whole BLA. The peak was more pronounced for young adults. Truncated power law provided poor fit for the distributions with R-squared being in the range 0.1 – 0.7 (except for the bin size < 4 ms in juvenile animals with R-square 0.9 and higher, Figure S2). Analysis of mean temporal profiles of amygdaloid avalanches with 2 SD threshold for the same randomly picked animals as in the previous section failed to reveal shape collapse (Figure S4B, S4D) in both age groups.

**Figure 3.**
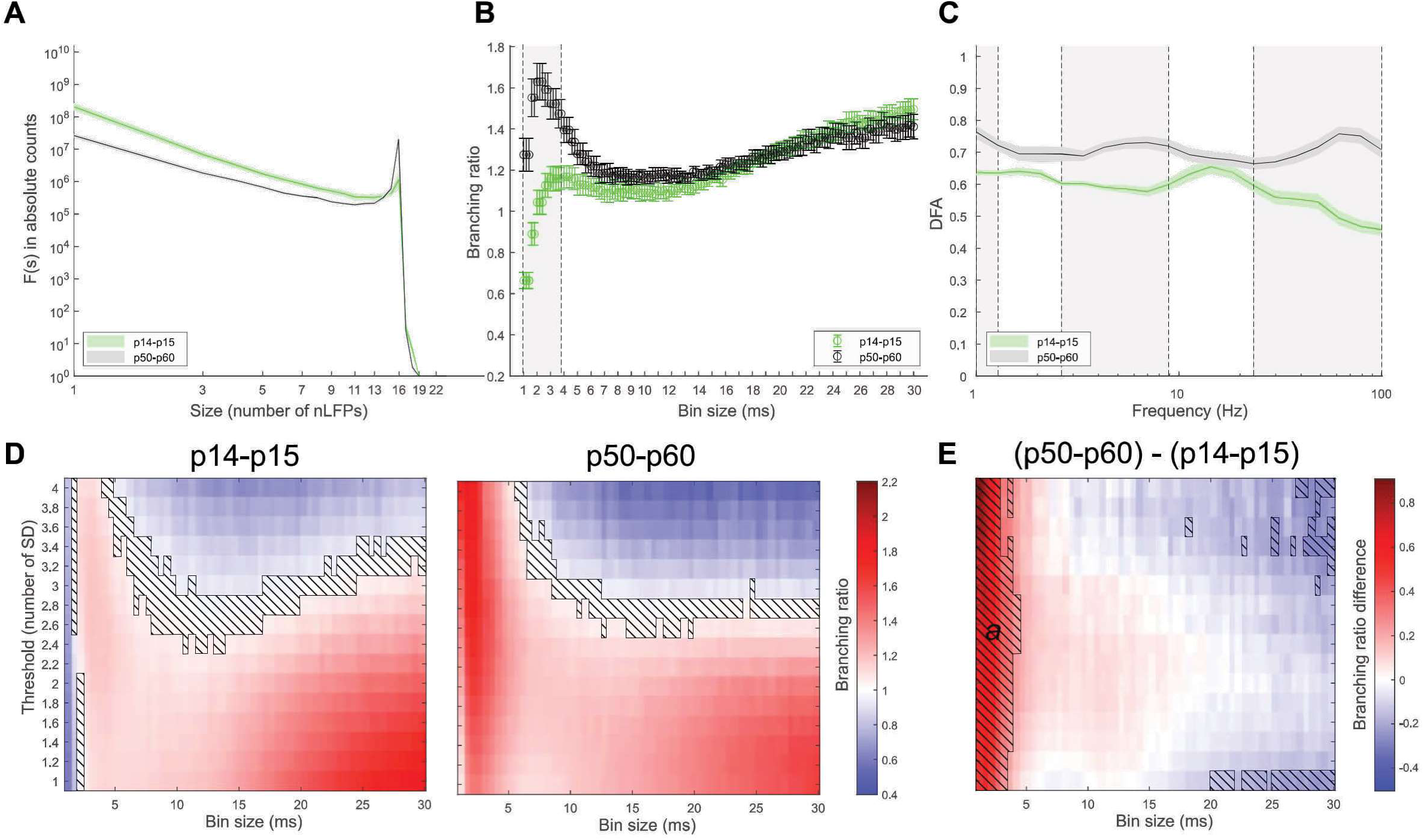
Neuronal avalanches and LRTCs across development in the BLA in male rats. (A) Avalanche size distribution calculated with 2SD threshold and 2 ms time bin and averaged across animals (N = 11, 13 for juvenile and adult animals, respectively). Size distribution deviates from the power law and is characterized by a sharp peak at 16 channels, more pronounced for young adult animals. (B) Mean branching ratios calculated with 2SD threshold for multiple time bins and averaged across animals (N = 11, 13 for juvenile and adult animals, respectively). (C) Mean DFA exponents calculated for 5 nearby channels and averaged across animals (N = 15, 18 for juvenile and adult animals, respectively). In (A-C) results for juvenile and adult animals are shown in green and black colors, respectively. Shaded area and error bars show SEM. Grey rectangles correspond to the significant differences (p < 0.05, Wilcoxon rank sum test with FDR correction, q < 0.05). (D) Mean branching ratios calculated with multiple thresholds and time bins and averaged across animals for juvenile (left panel) pups and young adult (right panel) rats. Hatched area corresponds to the branching ratios close to 1 (0.95 – 1.05). (E) Difference in the mean branching ratio between young adult and juvenile animals. Hatched areas correspond to the statistically significant differences (p < 0.05, Wilcoxon rank sum test with FDR correction, q < 0.05). Significant cluster ***a***: size s = 227, mean probabilistic index <*P(adult > juvenile)*> = 0.91.

Similar to the mPFC, branching ratio increased with the bin size (Figure 3B, 3D, S5B, S5D), however, there was a local maximum for small bin sizes (< 5 ms), more pronounced for adult animals. Moreover, for most parameters except large values of threshold (> 2.4 SD) branching ratio exceeded one. Such behavior of branching ratio has never been shown before for the brain. Taken together, peak in avalanche size distributions and high branching ratio show deviation from the critical state towards supercriticality.

In males, a comparison of two age groups revealed a significant increase in the mean branching ratio for the small bin sizes < 5 ms (Figure 3E, cluster ***a*,** p < 0.05, n = 11, 13, for juvenile and adult animals, respectively, Wilcoxon rank sum test with FDR correction, <*P(adult>juvenile)>* 0.91). Additionally, there were 12 small clusters of parameters with significant changes in the branching ratio. Similar increase for the small bin sizes < 3 ms is seen in female data (Figure S5E).

#### 3.2. Long-range temporal correlations in the BLA are present already in juvenescence and further increase over development

DFA analysis showed that in contrast to the mPFC, in the BLA positive LRTCs emerged already in juvenile animals (Figure 3C, S5C) where up to 30 Hz DFA exponents were close to 0.6, and increased in early adulthood with DFA exponents at the level of around 0.7 and higher for the whole frequency range (significant increase in all frequencies except 1.6 – 2 Hz and 11 – 18 Hz, p < 0.05, n = 15, 18, for juvenile and adult males, respectively, Wilcoxon rank sum test with FDR correction).

To sum up, in both age groups nLFPs in the BLA were clustered together into neuronal avalanches, however, avalanche size distributions could not be described by power law but instead were characterized by a sharp peak corresponding to the maximum number of channels, showing that once started an avalanche would likely extend to the whole system. Together with the branching ratio higher than one, this points towards the BLA operating in the supercritical regime. There was a developmental increase in the mean branching ratio for small bin sizes. Positive LRTCs in the amygdala emerged already in juvenescence and increased in adult animals. No sex-related differences were revealed in both avalanche characteristics and DFA exponents.

### 4. Cross-regional spread of avalanches is reduced over development

Intrigued by the unexpected behavior of the branching ratio in the BLA we decided to investigate interregional differences further. In both age groups there was a large cluster of parameters (cluster **a**, Figure 4A, S6A) where branching ratio was significantly higher in the BLA compared to the mPFC (p < 0.05, Wilcoxon signed-rank test with FDR correction, n = 11 in both juvenile and adult males) and this difference increased in adult animals. In juvenile males, cluster **a** comprised 728 points with mean probabilistic index <*P(BLA>mPFC)>* = 0.91. In adult males, cluster **a** comprised 936 points with <*P(BLA>mPFC)>* = 0.94. Similar results were seen in females though the effect was smaller (Figure S6A). Interestingly, the difference in the branching ratio between the regions disappeared with high values of the bin size. For instance, with 2 SD threshold there was no interregional difference in the branching ratio with bin sizes higher than 10 ms in juvenile males and higher than 15 ms in adult males (Figure 4B). Both regions have anatomical projections to one another with a conduction delay from BLA to the mPFC around 16 ms and from the mPFC to the BLA around 20 ms in adult rats (McGinty and Grace, 2008), which should be lower in smaller juvenile brain. In other words, significant differences in the branching ratio were restricted by the bin sizes less than a conduction delay, so activity from one region was not affecting another one.

**Figure 4.**
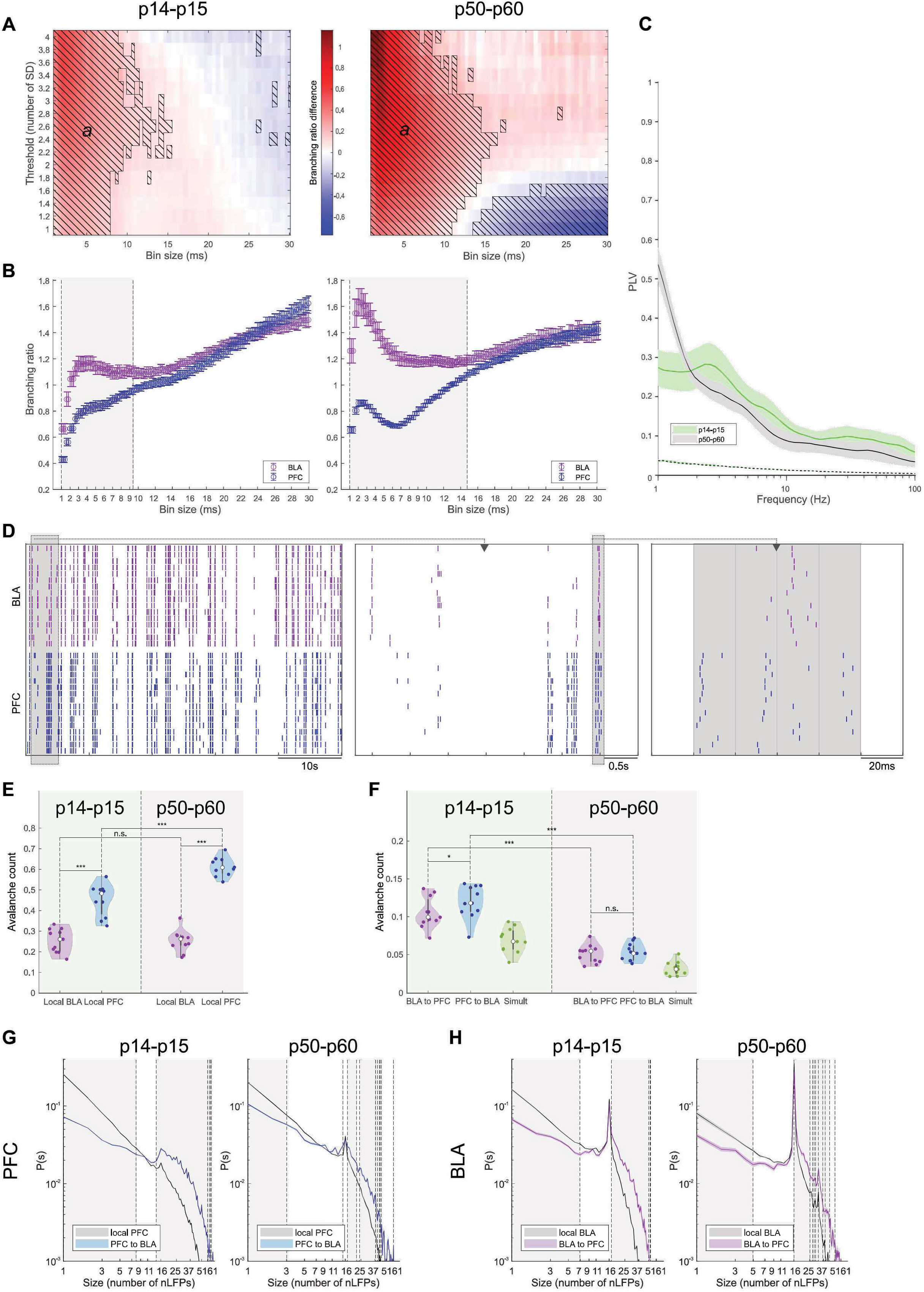
Spread of avalanches between the BLA and the mPFC in male rats. N = 11 in both juvenile and young adult group, if not stated otherwise. (A) Difference in the mean branching ratio of the avalanches in the BLA and the mPFC calculated for multiple thresholds and bin sizes. The mean branching ratio for the avalanches in the BLA is higher than in the mPFC for the small bin sizes. Hatched area corresponds to the statistically significant differences (p < 0.05, Wilcoxon signed-rank test with FDR correction, q < 0.05). Significant cluster ***a***: size s = 728, 936, mean probabilistic index *<P(BLA > PFC)>* = 0.91, 0.94 for juvenile and young adult animals, respectively. (B) Mean branching ratio averaged across animals (2SD threshold) for the avalanches in the mPFC (blue) and the BLA (magenta). Error bars show SEM. Grey rectangles correspond to significant differences (p < 0.05, Wilcoxon rank sum test with FDR correction, q < 0.05). In (A, B) juvenile and young adult data is shown on the left and right panel, respectively. (C) PLV between activity in the mPFC and the BLA for juvenile (in green) and young adult (in black) animals. Dashed line shows 95% confidence interval based on time-shuffled data. Shaded area corresponds to SEM (N = 14, 21 for juvenile and young adult animals, respectively). (D) An example of nLFPs (2SD threshold) detected simultaneously in the mPFC (blue) and the BLA (magenta) for a young adult rat. Panels represent different time scales with the largest one on the left. nLFP clusters can occur locally in one region or simultaneously in both regions. On the right panel, grey color highlights an avalanche starting in the mPFC and then emerging in the BLA (PFC to BLA avalanche). Time bin 20 ms. (E) Avalanche counts normalized by the total number of avalanches for local avalanches (2SD threshold, 20 ms bin size). Juvenile data is on the left panel and young adult data is on the right. Main effects cannot be properly interpreted because of the statistically significant interaction between avalanche localization and age group (age: F = 94.635, P < 0.001; localization: F = 137.933, P < 0.001; interaction: F = 11.432, P = 0.003). Post-hoc all pairwise multiple comparison: avalanche count is significantly higher in the mPFC than in the BLA in both age groups (juvenile: t = 5.914, p < 0.001; young adult: t = 10.695, p < 0.001); there is a developmental increase in avalanche count in the mPFC (t = 6.361, p < 0.001) but not in the BLA (t = 0.0339, p = 0.973). Two-way repeated measures ANOVA, Holm-Sidak method for all pairwise multiple comparisons. (F) Same as in (E) for two-region avalanches. Main effects: avalanche count is significantly higher for the avalanches starting in the mPFC compared to those starting in the BLA (F = 6.232, P = 0.021); there is a significant developmental decrease in the avalanche count for two-region avalanches (F = 88.848, P < 0.001); no interaction between avalanche directionality and age group (F = 1.249, P = 0.277). Post-hoc all pairwise multiple comparison: avalanche count is significantly higher for avalanches starting in the mPFC than those starting in the BLA in juvenile (t = 2.556, p = 0.019) but not in young adult (t = 0.975, p = 0.341) animals; there is a developmental decrease for both avalanches starting in the BLA (t = 8.131, p < 0.001) and in the mPFC (t = 9.051, p < 0.001). Two-way repeated measures ANOVA on log-transformed data, Holm-Sidak method for all pairwise multiple comparisons. (G) Normalized size distribution, calculated for local avalanches in the mPFC (*black*) and nLFP clusters in the mPFC that are followed by nLFP clusters in the BLA (*blue*) for juvenile (*left panel*) and young adult (*right panel*) males, averaged across animals. Shaded area corresponds to SEM. Grey rectangles and dashed vertical lines correspond to significant differences (p < 0.05, Wilcoxon signed-rank test with FDR correction, q < 0.05). NLFP clusters that are part of PFC to BLA avalanches are characterized by a reduced likelihood of small avalanches and an increased likelihood of large avalanches compared to the local PFC avalanches. (H) Same as (G) for local avalanches in the BLA (*black*) and nLFP clusters in the BLA that are followed by nLFP clusters in the mPFC (*magenta*).

We estimated functional connectivity between the regions. For both age groups, calculated PLV between the mPFC and the BLA was higher than expected by chance (Figure 4C, dashed line represents significance level) showing that there was a significant functional connectivity between the regions. Given anatomical and functional connectivity, we questioned whether neuronal avalanches in one region could trigger avalanches in another. Indeed, as can be seen from a raster plot of nLFPs (Figure 4D), there were clusters of nLFPs occurring in both regions simultaneously, which we would call two-region avalanches.

To delve deeper into this question, we calculated avalanches with one set of parameters. We picked a threshold of 2 SD, which provided a sufficient amount of nLFPs without picking up too much noise, and a bin size of 20 ms, time comparable with a conduction delay between two regions for adult rats. Then we estimated numbers of local (Figure 4 E, S6C) and two-region avalanches (Figure 4F, S6D).

For both age groups, most of the avalanches occurred locally either in the mPFC or the BLA (median (IQR) values 26.1% (10.5%) in the BLA and 48.3% (12%) in the mPFC in juvenile males, 26.2% (4.5%) in the BLA and 60.9% (6.4%) in the mPFC in young adult males, here and below n = 11 in both age groups). For local avalanches, two-way repeated measures ANOVA revealed significant interaction between age group and region (F = 11.432, P = 0.003) and thus main effects cannot be interpreted properly. Post-hoc all pairwise multiple comparison, Holm-Sidak method, showed a developmental increase in the relative number of local avalanches in the mPFC (t = 6.361, p < 0.001) but not in the BLA (t = 0.0339, p = 0.973). For both age groups relative number of avalanches in the mPFC was higher than in the BLA (juvenile: t = 5.914, p < 0.001; young adult: t = 10.695, p < 0.001).

A minor number of avalanches spanned both regions. Here we would say that an nLFP cluster A from one region is following an nLFP cluster B from the second region if it starts in any time bin in the range t_2_ – t_end+1_, where t_2_ is the second and t_end_ is the last bin of the cluster B. Both clusters A and B would be counted as one avalanche. From two-region avalanches the smallest group (6.7% (2.5%) in juvenile and 3.1% (1.2%) in adult males) were avalanches starting simultaneously in both regions (time delay between the first nLFP in each region was less than one time bin). However, most of the two-region avalanches started with nLFPs in one region and were followed by nLFPs in the second one (in juvenile males: median (IQR) values 11.8% (3.3%) and 9.9% (2.95%) for avalanches starting in the mPFC (PFC to BLA avalanches) and the BLA (BLA to PFC avalanches), respectively; in young adult males: 5.4% (1.6%) and 5.1% (1.9%)). Two-way repeated measures ANOVA on log-transformed data (raw data did not pass equal variance assumption, p < 0.05, Levene’s test) revealed no significant interaction between age group and avalanche directionality (F = 1.249, P = 0.277). Significant differences were found between different levels of both age and directionality after allowing for the effects of differences in the second factor. There was a significant developmental decrease in the avalanche count for two-region avalanches (F = 88.848, P < 0.001); an avalanche count was significantly higher for the avalanches starting in the mPFC compared to those starting in the BLA (F = 6.232, P = 0.021). Post-hoc all pairwise multiple comparison, Holm-Sidak method, showed a developmental decrease in the relative number of two-region avalanches (PFC to BLA: t = 9.051, p < 0.001; BLA to PFC: t = 8.131, p < 0.001). The relative number of PFC to BLA avalanches was higher compared to those starting in the BLA in juvenile (t = 2.556, p = 0.019), but not in adult group (t = 0.975, p = 0.341). An estimation of avalanche directionality in females gave similar results (Figure S6).

Next, we asked if there is a size difference between nLFP clusters that stay local and those that spread to another region. Figure 4G depicts normalized size distributions calculated separately for local PFC avalanches and nLFP clusters in the PFC that are part of PFC to BLA avalanches. The latter was characterized by the reduced probability of small cluster sizes (s < 8 and s < 3 for juvenile and young adult males, respectively) and increased probability of large cluster sizes (s > 14 and s > 17 for juvenile and young adult males, respectively, p < 0.05, Wilcoxon signed-rank test with FDR correction). Reduced probability of small cluster sizes (s < 7 and s < 5 for juvenile and young adult males, respectively) and increased probability of large cluster sizes (s > 17 and s > 16 for juvenile and young adult males, respectively, p < 0.05, Wilcoxon signed-rank test with FDR correction) was also observed in the BLA for the nLFP clusters that spread to the mPFC compared to those that stay local (Figure 4H). The same was found in females (Figure S6E, S6F). In other words, nLFP clusters spreading from one region to another tend to be larger than those that stay local. Note, however, that here we use the terms “spread” and “stay local” in a descriptive manner without claiming causality. We cannot say if nLFP clusters in one region are caused by nLFP clusters in another region or if they occur independently at the same time, as well as we cannot exclude interaction with any other region.

To sum up, in both age groups most avalanches were localized in one region, with more avalanches occurring in the mPFC. However, a small proportion of avalanches spanned both regions. From them, there was a slightly higher number of avalanches starting in the mPFC compared to those starting in the BLA in juvenile animals, whereas in young adults the leading role could not be attributed to one region or another. With age we saw an increase in the relative number of local avalanches in the mPFC accompanied by a decrease in the number of two-region avalanches. Noteworthy, in both age groups and in both regions nLFP clusters that spread to another region tended to be larger than those that stayed local.

## Discussion

Extensive in vitro and in vivo evidence indicates that critical neuronal dynamics and self-organized criticality are intimately involved in the development of neuronal networks (Gireesh and Plenz, 2008). In this study, we focused on differences in critical dynamics between the mPFC and BLA, two closely interconnected brain regions. The criticality hypothesis suggests that neural networks naturally self-organize toward a transitional state—the critical point—situated between an ordered (subcritical) regime, where activity rapidly dissipates, and a disordered (supercritical) regime, where activity escalates uncontrollably. At this state, the network is thought to achieve maximal dynamic range (Kinouchi and Copelli, 2006; Shew *et al*., 2009), information-processing capacity (Beggs and Plenz, 2003; Haldeman and Beggs, 2005; Shew *et al*., 2011), and complexity (Timme *et al*., 2016). In cognitive processes, mPFC is thought to participate in large-scale hierarchical processing required in attention, memory, and decision-making (Dalley, Cardinal and Robbins, 2004; Lara and Wallis, 2015; Friedman and Robbins, 2022) and its functional anatomy is characterized by highly laminar structure and dense recurrent excitatory and inhibitory connectivity (Petrides, 2005; Anastasiades and Carter, 2021). In contrast, the BLA is specialized for the rapid integration of sensory inputs with limbic and autonomic outputs. The BLA operates as a fast-acting detector and regulator specialized for emotional significance, fear learning, and survival behaviors (Phelps and LeDoux, 2005; Janak and Tye, 2015), characterized by more tightly coupled, stimulus-driven dynamics. The amygdala exhibits a nuclei-based organization (Sah *et al*., 2003; McDonald, 2020), unlike the layered structure of the mPFC, and its activity is burst-like, tightly linked to salient stimuli such as threats and rewards (Wassum, 2022).

Neuronal avalanches can be seen in the acute brain slices (Beggs and Plenz, 2003), organotypic (Stewart and Plenz, 2006, 2008; Plenz *et al*., 2011) and dissociated neuronal cultures (Pasquale *et al*., 2008; Yada *et al*., 2017) *in vitro*, in the animal *in vivo* LFP recordings (Gireesh and Plenz, 2008; Petermann *et al*., 2009; Priesemann, Munk and Wibral, 2009), as well as in large-scale recording techniques such as MEG (Palva *et al*., 2013; Shriki *et al*., 2013; Zhigalov *et al*., 2015; Lombardi *et al*., 2023), EEG (Palva *et al*., 2013; Lombardi *et al*., 2023; Scarpetta *et al*., 2023; Maschke *et al*., 2024), and fMRI (Xu, Feng and Yu, 2022; Xin *et al*., 2025) in human brain. The power-law distribution of avalanche sizes suggests that these events span multiple orders of magnitude, occurring across all scales including those large enough to engage activity in multiple, distinct brain regions. These findings are in line with the idea that neuronal avalanches facilitate long-range communication within the brain. However, it must be considered that functional connectivity is shaped by the underlying structural connectivity and that the two are strongly interrelated (Lim *et al*., 2019). Therefore, we focused on uncovering the details of local critical dynamics in the mPFC–BLA circuitry.

### Signal characteristics in mPFC and BLA

Typical for urethane anesthesia, sleep-like slow-wave delta activity was predominant in both juvenile and young adult animals, the latter group displayed brain-state alterations like those described in (Clement *et al*., 2008; Pagliardini *et al*., 2012; Silver, Ward-Flanagan and Dickson, 2021). With development, a peak in the delta power shifted towards faster frequencies and there was a developmental increase in theta- and gamma-power. This is in line with developmental power alterations described in the BLA and the mPFC under urethane anesthesia in mice (Donati, Vedele and Hartung, 2024) and in the PFC of awake mice (Pöpplau *et al*., 2024). Both regions were characterized by high inter-regional functional connectivity, increasing with age.

### LRTCs

An important property of the critical systems is maximized memory capacity (Haldeman and Beggs, 2005), which is reflected in high long-range temporal correlations (LRTCs), whereas deviations from criticality to both sub- and supercritical dynamics are associated with decrease in LRTCs (Poil *et al*., 2012). As the emergence of LRTCs depends on excitation/inhibition balance (Poil *et al*., 2012), we asked how developmental changes in the prefrontal-amygdala network affect it. To reveal this, we exploited detrended fluctuation analysis (DFA), introduced by (Peng *et al*., 1994).

mPFC: In the juveniles DFA exponents in mPFC were close to 0.5 thus suggesting absence of LRTCs. Positive LRTCs emerged in PFC only in young adult animals, where DFA exponents were significantly higher compared to juvenile animals for the whole frequency range. LRTCs are essential for prefrontal functions, such as attention and working memory (Meisel *et al*., 2013; Simola *et al*., 2017), and thus our results highlight functional immaturity of the juvenile mPFC and are in line with prolonged functional prefrontal development (Pöpplau and Hanganu-Opatz, 2024).BLA: In BLA positive LRTCs emerged already in juvenile animals where up to 30 Hz DFA exponents were close to 0.6 and increased in early adulthood with DFA exponents at the level of around 0.7 for the whole frequency range. In keeping with the central role of the BLA in sustaining immediate vital behaviors, strong LRTCs in the BLA could represent one of the neuronal adaptations that optimize amygdaloid circuitry for responding to sustained physiological challenges.

Our results are in line with a prolonged, spanning into young adulthood, augmentation of long-range temporal correlations reported in humans (Smit *et al*., 2011), and reflect maturation of the brain. It is important, however, to consider that the developmental increase in the DFA exponents in our data in both regions coincided with an increase in oscillatory power in a broad band and as such can be partially explained by increased signal-to-noise ratio as stressed in (Linkenkaer-Hansen *et al*., 2007).

### Branching ratios

A branching ratio is defined as the mean number of downstream neurons activated by a single active neuron during an avalanche, thus quantifying the propagation of activity through the network (Kinouchi and Prado, 1999).

mPFC: In general, branching ratios in the juvenile mPFC were higher than those in the adult mPFC, reflecting more promiscuous neuronal activity in the younger cortex. These findings are in line with the known alterations in synaptic density and synaptic pruning, neuronal apoptosis, as well as interneuronal maturation and the formation of perineuronal nets that occur during mPFC development (reviewed in (Zimmermann, Richardson and Baker, 2019; Juraska and Drzewiecki, 2020; Pöpplau and Hanganu-Opatz, 2024)). Interestingly, while the degree of the postpubertal non-GABAergic neuronal loss was shown to be considerably larger in female rats (Markham, Morris and Juraska, 2007; Willing and Juraska, 2015) and dendritic retraction was seen only in females (Koss *et al*., 2014), the decrease in the branching ratio in our results was significant only in males indicating more profound changes in the male prefrontal network dynamics.

BLA: For both age groups, amygdala avalanches were characterized by a high mean branching ratio exceeding one for most of the parameters. In terms of development, in general the mean branching ratio in the BLA stayed stable over time, highlighting a divergence from the prefrontal developmental trajectory. Indeed, though some postpubertal neuronal and glial apoptosis takes place in the BLA, amygdala volume increases over time (Rubinow and Juraska, 2009) and, unlike the PFC, the amygdala does not undergo synaptic pruning but rather is characterized by an increasing arborization and spine density which stabilizes after adolescence (Koss *et al*., 2014).

It is worth mentioning that in both mPFC and BLA we see a developmental increase in the branching ratio for very small bin sizes, however, the timescales less than propagation delay (Madadi Asl, Valizadeh and Tass, 2018) are not sufficient to reflect spread of activity between the neurons but rather delays in the processing of the common input. Hence, this increase is likely related to the developmental changes in the external drive or sensitivity to it.

### Power laws and shape collapse

mPFC: As was previously shown for the cortical avalanches (Beggs and Plenz, 2003; Gireesh and Plenz, 2008) in both age groups size distributions of prefrontal avalanches followed truncated power law as expected for the systems operating near a critical state. In case of young adults, we also showed shape collapse of the avalanche mean temporal profiles, another hallmark of critical systems described in neuronal networks (Friedman *et al*., 2012). In juvenile animals, avalanche shapes collapsed to a lower extent.

BLA: In stark contrast to the mPFC, avalanche size distributions in the BLA could not be described by power law and instead were characterized by a sharp peak at the maximum number of analyzed channels indicating that large number of avalanches span the whole BLA. Such avalanches represent so-called dragon king phenomenon (Sornette, 2009), events with characteristic scale coexisting with scale-free dynamics of smaller events, described in various systems and arising from positive feedback mechanisms. Interestingly, coexistence of small-sized power law avalanches and supercritical bumps in self-organizing neural networks was predicted in several mathematical models (Costa, Brochini and Kinouchi, 2017; Kinouchi *et al*., 2019) and our results provide first experimental evidence of this. We also failed to show shape collapse of the mean temporal profiles in the amygdala avalanches in both age groups. Taken together with a high branching ratio, this shows that, unlike the cortex, the BLA does not operate near critical state deviating towards supercriticality. While supercriticality is commonly associated with disease (Zimmern, 2020; Heiney *et al*., 2021), it can have functional relevance in a healthy state. For instance, Shew et al. (Shew *et al*., 2015) detected transient supercritical state in the turtle visual cortex during sensory adaptation to the movie onset; such shifts towards supercriticality allow strong response to important transients.

Since both PFC and BLA have roughly the same prevalence of GABAergic neurons (reviewed in (Hájos, 2021; Kupferschmidt *et al*., 2022)), the stark difference in regional dynamics cannot be explained by anatomical excitation/inhibition balance but likely arises from different anatomical organizations. Network topology indeed was shown to be crucial for maintaining criticality (Massobrio, Pasquale and Martinoia, 2015; Sugimoto, Yadohisa and Abe, 2025) as well as capable of giving rise to dragon king avalanches (Sugimoto, Yadohisa and Abe, 2025). Distinct dynamical states of the BLA and the PFC go in line with different functional demands. While optimized information processing achieved near critical state is crucial for prefrontal executive functions, the amygdala is directly involved in essential survival behaviors benefiting from strong response to possible threats which can be achieved by the amplified propagation of the signal.

### mPFC-BLA crosstalk

The BLA and the PFC are strongly reciprocally interconnected with the corresponding projections emerging by the end of the second development week (Bouwmeester, Smits and Van Ree, 2002; Bouwmeester, Wolterink and Van Ree, 2002) and undergoing intense anatomical and functional reorganization during adolescence. Projections from the BLA are established slightly earlier than the prefrontal ones and continue to proliferate until adulthood (Cunningham, Bhattacharyya and Benes, 2002) while prefrontal projections to the BLA peak during adolescence, then undergo pruning and stabilize at adulthood (Cressman *et al*., 2010). Given these, we asked if avalanches can be transferred between the regions and how this transfer is affected by projection development.

For both age groups, calculated PLV between mPFC and BLA shows that there was a significant functional connectivity between the regions. Regarding critical dynamics, in both age groups most avalanches were localized within a single region, with more avalanches occurring in the mPFC. However, a small proportion of avalanches spanned both regions. Among these, there was a slightly higher number of avalanches starting in the mPFC compared to those starting in the BLA in juvenile animals, whereas in young adults the leading role could not be attributed to either region. With age, we observed an increase in the relative number of local avalanches in the mPFC, accompanied by a decrease in the number of two-region avalanches. At first glance, this decrease contradicts data on the connectivity maturation between the regions. However, both amygdala and prefrontal projections innervate not only GLU-but also GABAergic neurons (Cunningham, Bhattacharyya and Benes, 2008; Hübner *et al*., 2014; McGarry and Carter, 2016). Interaction of the amygdala afferents with the prefrontal interneurons was shown to increase until adulthood (Cunningham, Bhattacharyya and Benes, 2008) and while prefrontally recruited excitation in the basal amygdala once established stays stable over time feedforward inhibition of the BA undergoes enhancement up until late adolescence (Arruda-Carvalho *et al*., 2017), which is also in line with the functional prefrontal-amygdala connectivity shift to negative valence (Gee *et al*., 2013) during adolescence. Developmental increase in feedforward inhibition recruited by prefrontal and amygdala projections can result in reduced avalanche transfer between the regions.

Notably, in both age groups and in both regions, nLFP clusters that spread to the other region tended to be larger than those that remained local, suggesting a filtering mechanism favoring transfer of large activity clusters. A more detailed study of excitation/feedforward inhibition interaction in the target regions can be a key to understanding this phenomenon.

### Study limitations

Since amygdala is highly sensitive to acute stress and timescales for recordings in juvenile animals do not allow for habituation, experiments were performed under urethane anaesthesia, which is a major limitation of the study. While urethane anaesthesia is shown to mimic normal sleep (Clement *et al*., 2008; Ward-Flanagan *et al*., 2022) and studies on anaesthetized animals (Gireesh and Plenz, 2008; Hahn *et al*., 2010) have shown power-law distributions of cortical avalanches, emergent body of evidence shows that vigilance state (Priesemann *et al*., 2013; Scarpetta *et al*., 2023) and anaesthesia induced changes in the brain state (Fagerholm *et al*., 2018; Curic *et al*., 2024; Maschke *et al*., 2024) alter avalanche characteristics. As such, we cannot rule out the possibility that our results are affected by anaesthesia. Our comparisons remain meaningful under the assumption that anaesthesia affects neuronal dynamics in both regions at both developmental time points similarly. In juvenile animals with discontinuous activity (before p 12) anaesthesia increases signal discontinuity and thus effects differ from adults (see (Chini and Hanganu-Opatz, 2021)); however, this should not be the case for continuous activity that we had in our recordings. Our second concern is regional differences in urethane sensitivity. If the cortex is more sensible to urethane anaesthesia than the BLA, reduced inhibition from the mPFC can shift excitation/inhibition balance in the amygdala and lead to the supercritical bump. Additional experiments on awake animals are needed to exclude this possibility.

### Conclusions

To conclude, we have described neuronal avalanches in the prefrontal-amygdala network in juvenile and young adult rats and shown developmental decrease in the mean branching ratio in the mPFC reflecting known prefrontal maturational changes taking place during adolescence. We have shown that the mPFC and the BLA operate in different dynamical regimes. While prefrontal avalanches exhibited scale-free properties of the size distributions at both developmental stages, avalanches in the amygdala tended to span the whole region favoring maximal size. Taken together with branching ratio exceeding one and the absence of shape collapse, this points towards supercritical regime of the amygdala network dynamics, providing evidence that while the brain is thought to operate near critical state, individual regions can benefit from other dynamical regimes. Additionally, we have shown that neuronal avalanches can span both regions; however, efficacy of avalanche transfer is reduced with the development of the prefrontal-amygdala network.

## Acknowledgments

This study was financially supported by Jane and Aatos Erkko Foundation and the Research Council of Finland, grant number 361235.

We thank Janne Sulku, Ehtasham Javed, the personnel in the Laboratory Animal Center in University of Helsinki for technical help.

## Conflict of interest statement

The authors declare no conflict of interest.

## Credit author statement

Z.K.: carried out the electrophysiological experiments and data analysis, contributed to the data analysis tools. M.P., S.P., S.L., H.H., T.T.: contributed to methodology design. H.H., T.T.: provided resources for the experimental work, conceptualized and supervised the project. The manuscript was written by Z.K. and T.T. with contribution of S.L. and H.H.

---

**Figure S1.**
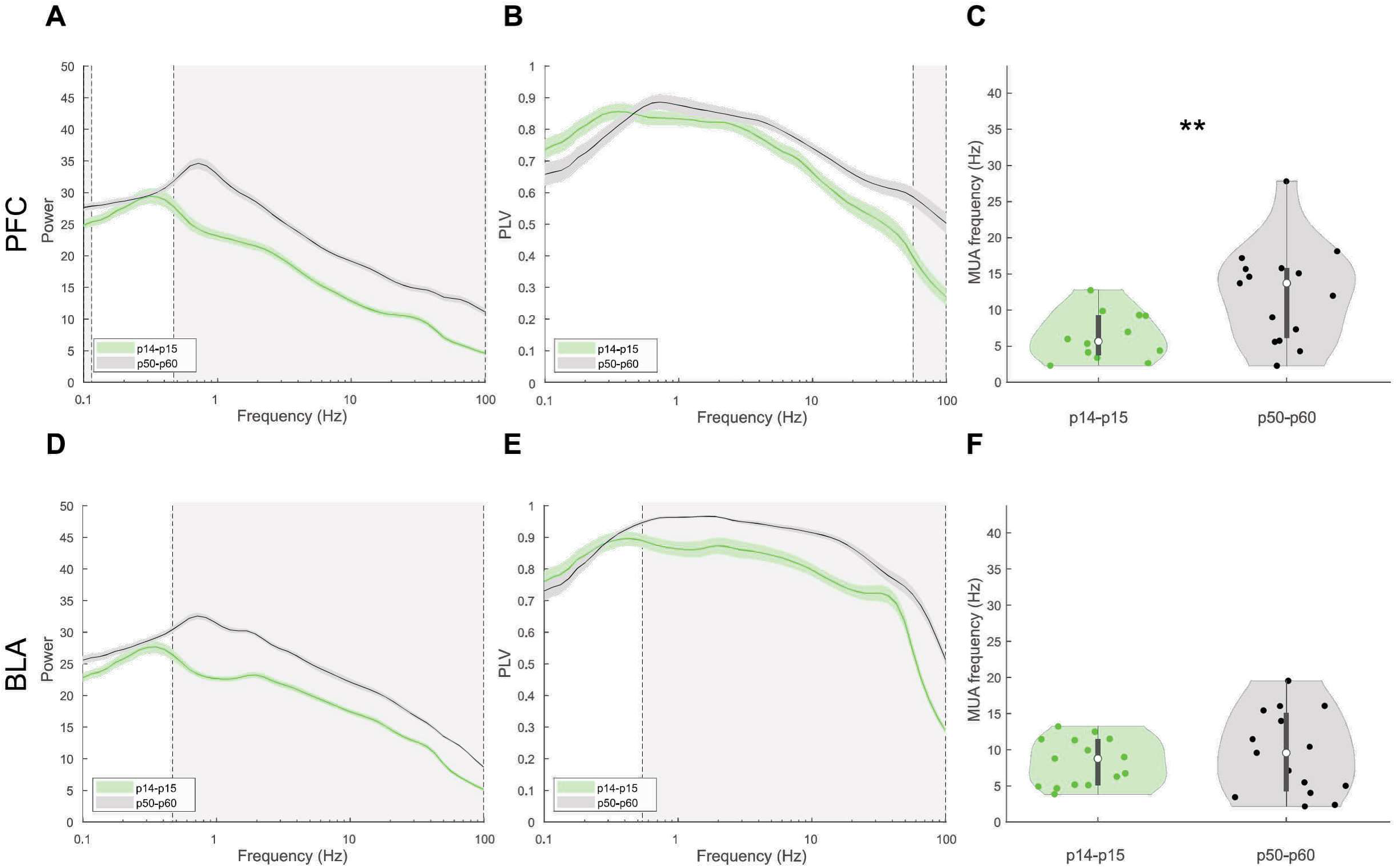
LFP signal in the mPFC and the BLA in female rats. (A, D) Mean power spectra calculated for 16 channels in the mPFC (A) and the BLA (D) from juvenile (green, N = 15, 17 for the mPFC and the BLA, respectively) and young adult (black, N= 15 animals for both mPFC and BLA) female rats. Shaded area shows SEM. Grey rectangles represent significant differences. (B, E) Same as (A, D) for PLV. (C, F) Instantaneous MUA firing rate (Hz) calculated for a single channel in the mPFC (C) and the BLA (F) of juvenile (green, N = 12, 15 for the mPFC and the BLA, respectively) and young adult (black, N = 15 for both mPFC and BLA) rats. Firing rates increase with age in the mPFC (juveniles: median (IQR) = 5.67 (5.51), young adults: median (IQR) = 13.72 (9.62), p = 0.007, Welch’s t-test), but remain similar across development in the BLA (juveniles: median (IQR) = 8.77 (6.32), young adults: median (IQR) = 9.56 (10.79), p = 0.494, Welch’s t-test).

**Figure S2.**
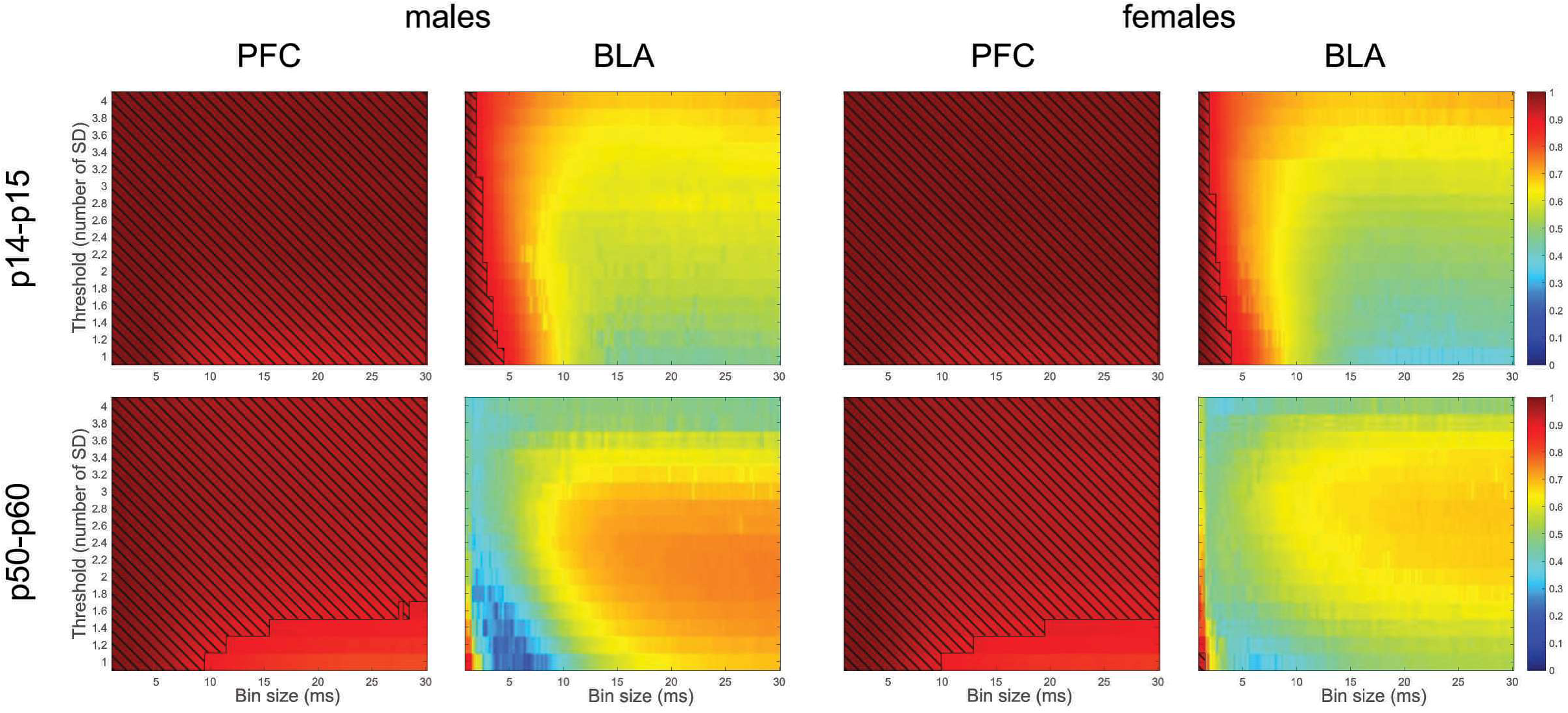
Truncated power-law goodness of fit (R-squared) for avalanche size distributions in juvenile (upper panels) and adult (lower panels) animals. Hatched areas represent values with R-squared higher than 0.9. In the mPFC but not in the BLA in both age groups for most of the parameters avalanche size distributions follow truncated power-law.

**Figure S3.**
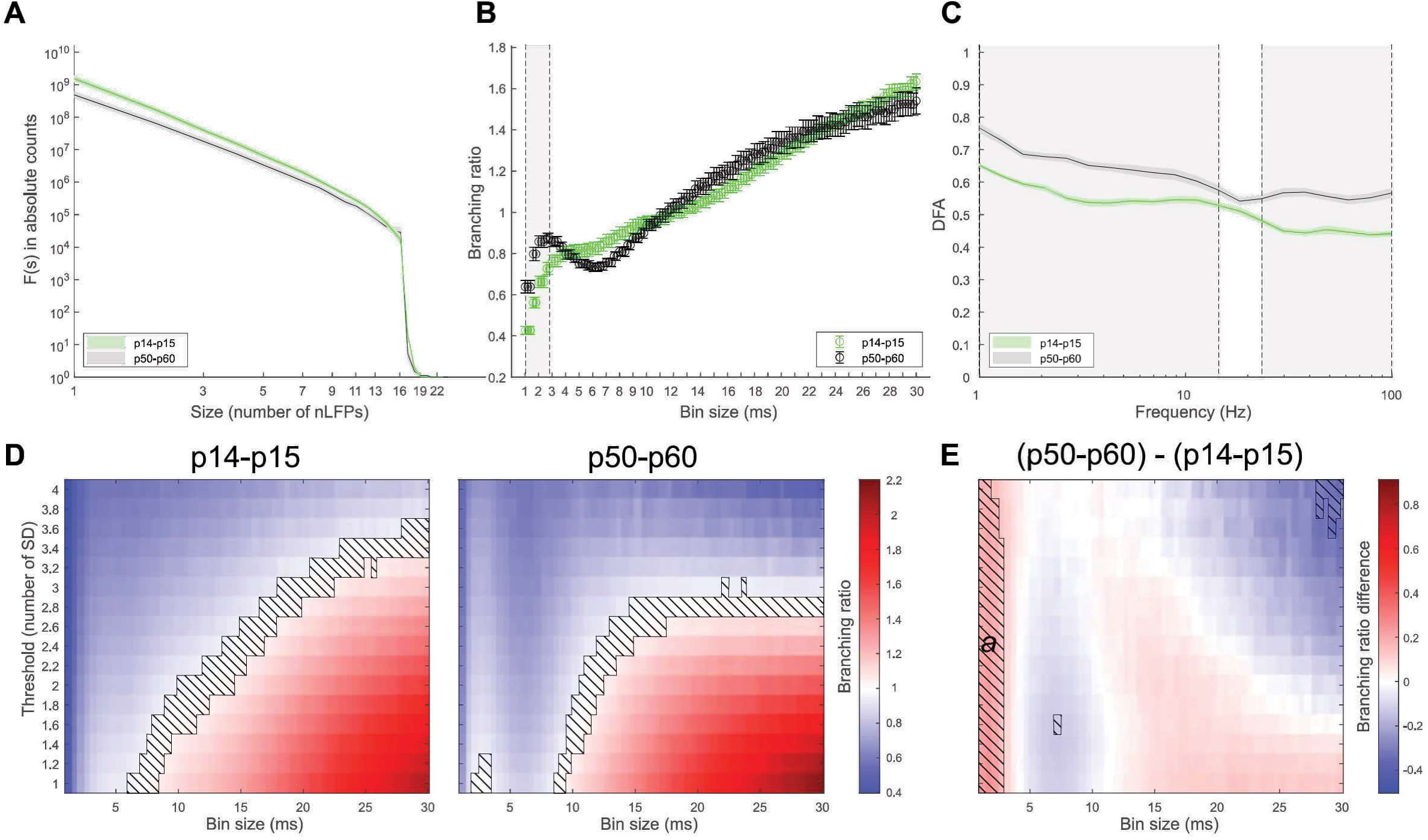
Neuronal avalanches and LRTCs across development in the mPFC in female rats. (A) Avalanche size distribution calculated with 2SD threshold and 2 ms time bin and averaged across animals (N= 15 for both juvenile and young adult animals). In log-log scale, distribution represents straight line a with cut-off at 16 channels and is described by truncated power law. (B) Branching ratios calculated with 2SD threshold for multiple time bins and averaged across animals (N = 15 for both juvenile and young adult animals). (C) Mean DFA exponents calculated for five nearby channels and averaged across animals (N = 15, 19 for juvenile and young adult animals, respectively). In *A-C* results for juvenile and young adult animals are shown in green and black colors, respectively. Shaded area and error bars show SEM. Grey rectangles correspond to the significant differences (p < 0.05, Wilcoxon rank sum test with FDR correction, q < 0.05). (D) Mean branching ratios calculated with multiple thresholds and time bins and averaged across animals for juvenile (left panel) pups and young adult (right panel) rats. Hatched area corresponds to the branching ratios close to 1 (0.95 – 1.05). (E) Difference in the mean branching ratio between young adult and juvenile animals. Hatched areas correspond to the statistically significant differences (p < 0.05, Wilcoxon rank sum test with FDR correction, q < 0.05). Significant cluster ***a***: size s = 151, mean probabilistic index *<P(adult > juvenile)>* = 0.91.

**Figure S4.**
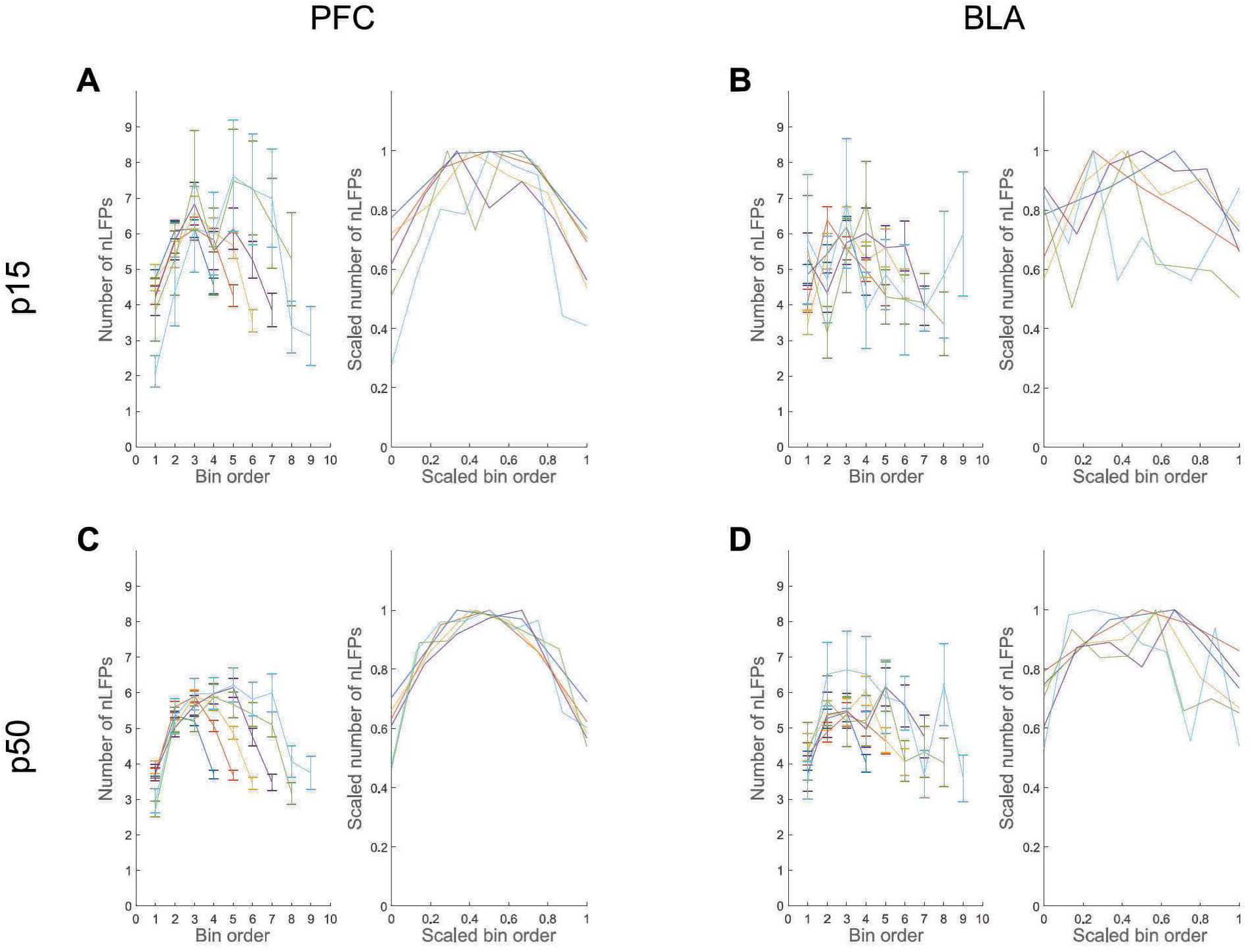
An example of raw (mean and SEM, *left panels*) and scaled (normalized mean, *right panels*) mean avalanche profile for an individual animal in the mPFC (A) and BLA (B) of a juvenile male rat (p15) calculated with 2 SD threshold and 16 ms time bin. Avalanche profiles are shown for the avalanches with durations 4 (*blue curve*), 5 (*red curve*), 6 (*yellow curve*), 7 (*purple curve*), 8 (*green curve*) and 9 (*light blue*) bins. (C, D) Same as (A, B) for young adult male rat (p50). Mean avalanche profiles in the mPFC of young adult rat are bell-shaped and, when scaled, display shape collapse.

**Figure S5.**
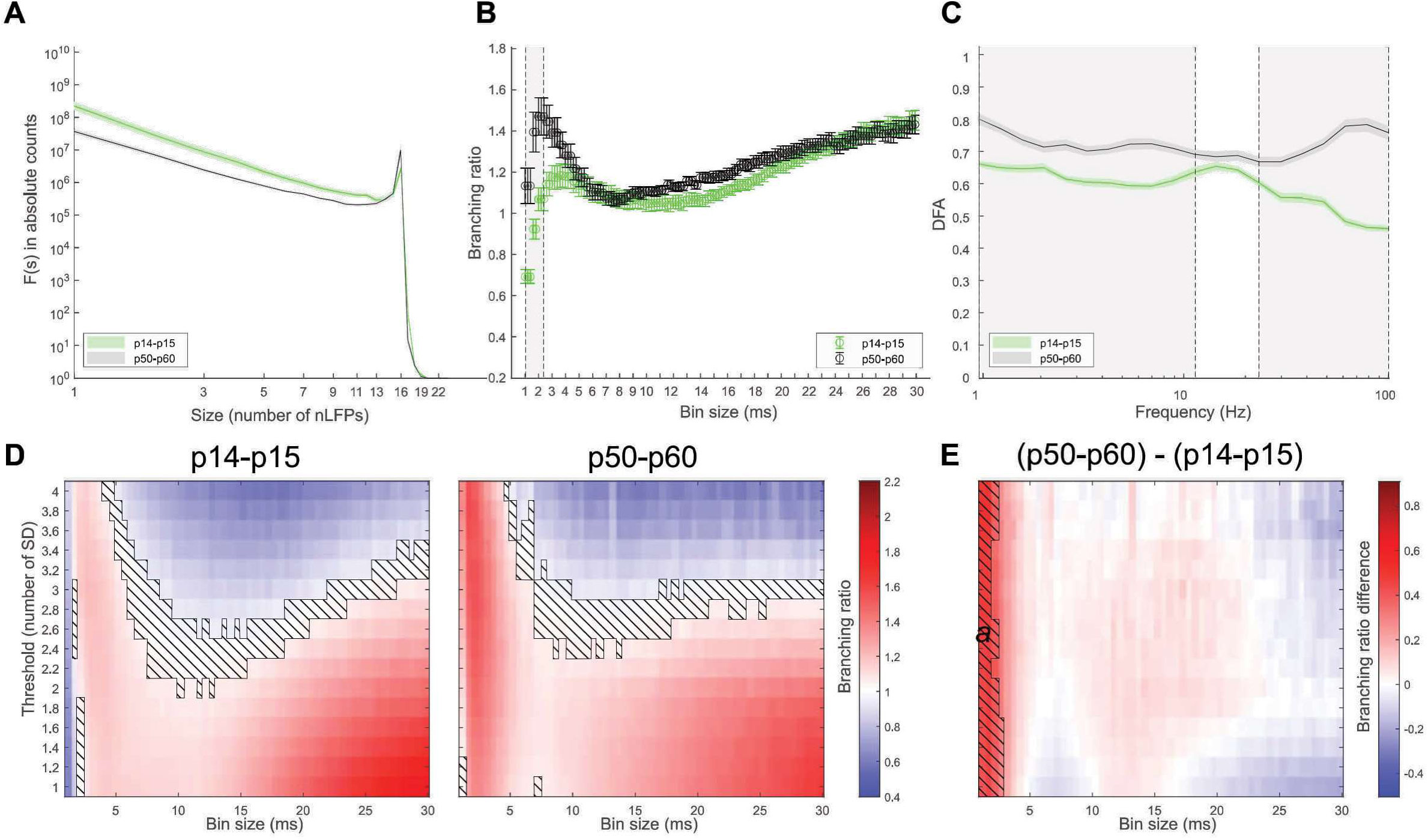
Neuronal avalanches and LRTCs across development in the BLA in female rats. (A) Avalanche size distribution calculated with 2SD threshold and 2 ms time bin and averaged across animals (N = 17, 15 for juvenile and young adult animals, respectively). Size distribution deviates from power law and is characterized by a sharp peak at 16 channels, more pronounced for young adult animals. (B) Mean branching ratios calculated with 2SD threshold for multiple time bins and averaged across animals (N = 17, 15 for juvenile and young adult animals, respectively). (C) Mean DFA exponents calculated for 5 nearby channels and averaged across animals (N = 18 for both juvenile and young adult animals). In (A-C) results for juvenile and young adult animals are shown in green and black colors, respectively. Shaded area and error bars show SEM. Grey rectangles correspond to significant differences (p < 0.05, Wilcoxon rank sum test with FDR correction, q < 0.05). (D) Mean branching ratios calculated with multiple thresholds and time bins and averaged across animals for juvenile (left panel) pups and young adult (right panel) rats. Hatched area corresponds to the branching ratios close to 1 (0.95 – 1.05). (E) Difference in the mean branching ratio between adult and juvenile animals. Hatched areas correspond to the statistically significant differences (p < 0.05, Wilcoxon rank sum test with FDR correction, q < 0.05). Significant cluster ***a***: size s = 121, mean probabilistic index *<P(adult > juvenile)>* = 0.84.

**Figure S6.**
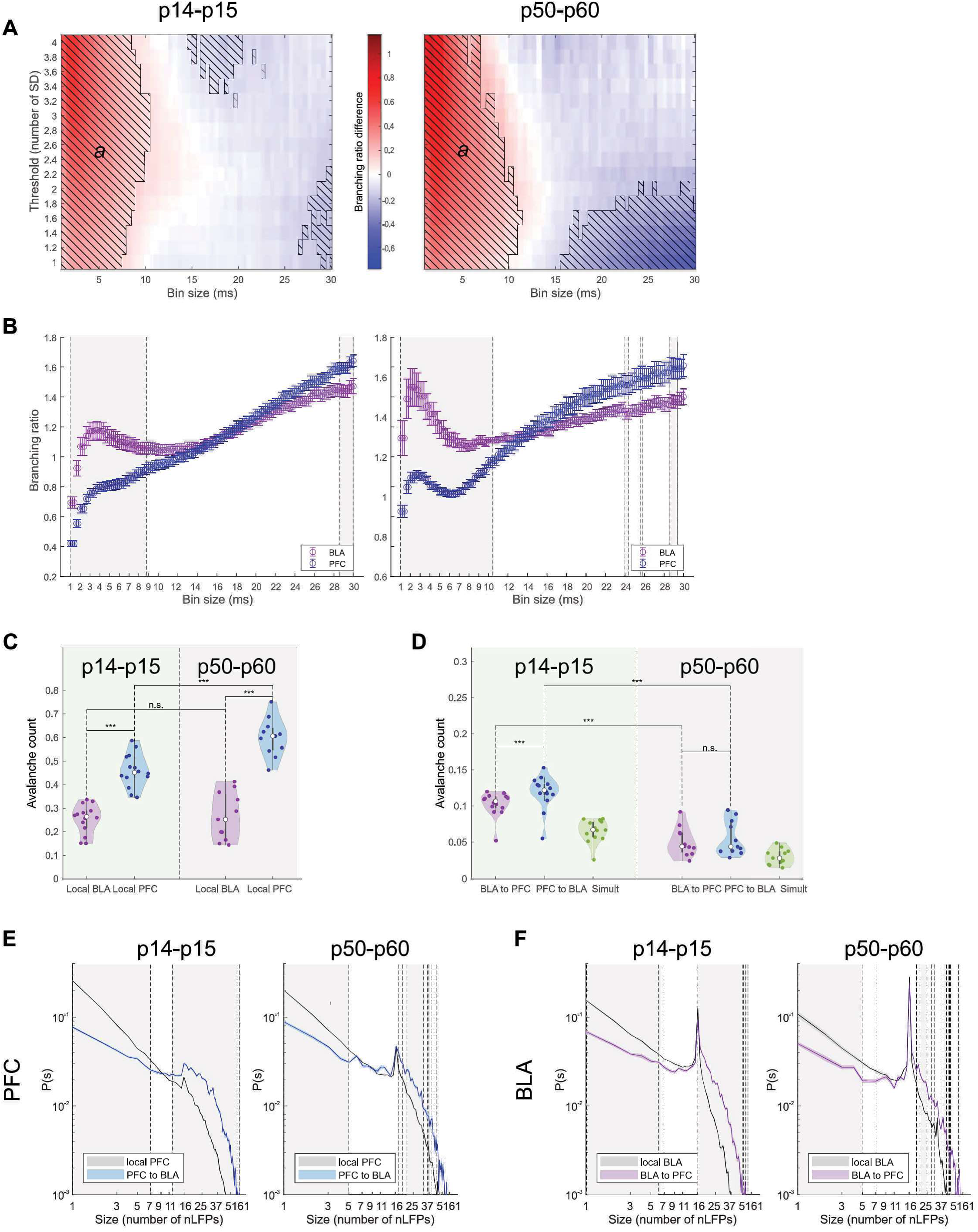
Spread of avalanches between the BLA and the mPFC in female rats. N = 14 and 11 in juvenile and young adult group, respectively. (A) Difference in the mean branching ratio of avalanches in the BLA and the mPFC calculated for multiple thresholds and bin sizes. The mean branching ratio for the avalanches in the BLA is higher than in the mPFC for the small bin sizes. Hatched area corresponds to the statistically significant differences (Wilcoxon signed-rank test with FDR correction, q < 0.05). Significant cluster ***a***: size s = 667, 657, mean probabilistic index: *<P(BLA > PFC)>* = 0.89, 0.92 for juvenile and young adult animals, respectively. (B) Mean branching ratio averaged across animals (2SD threshold) in the mPFC (blue) and the BLA (magenta). Error bars show SEM. Grey rectangles correspond to the significant differences (p < 0.05, Wilcoxon rank sum test with FDR correction, q < 0.05). In (A, B) juvenile and young adult data is shown on the left and right panel, respectively. (C) Avalanche counts normalized by the total number of avalanches (2SD threshold, 20 ms bin size) for local avalanches. Juvenile data is on the left panel and young adult data is on the right. Main effects cannot be properly interpreted because of the statistically significant interaction between avalanche localization and age group (age: F = 53.317, P < 0.001; localization: F = 83.017, P < 0.001; interaction: F = 4.828, P = 0.038). Post-hoc all pairwise multiple comparison: avalanche count is significantly higher in the mPFC than in the BLA in both age groups (juvenile: t = 5.212, p < 0.001; young adult: t = 7.556, p < 0.001); there is a developmental increase in the avalanche count in the mPFC (t = 4.480, p < 0.001) but not in the BLA (t = 0.331, p = 0.743). Two-way repeated measures ANOVA, Holm-Sidak method for all pairwise multiple comparisons. (D) Same as in (C) for two-region avalanches. Main effects: the avalanche count is significantly higher for the avalanches starting in the mPFC compared to those starting in the BLA (F = 12.764, P = 0.002); there is a significant developmental decrease in the avalanche count for two-region avalanches (F = 51.586, P < 0.001); no interaction between avalanche directionality and age group (F = 3.796, P = 0.064). Post-hoc all pairwise multiple comparison: avalanche count is significantly higher for the avalanches starting in the mPFC than those starting in the BLA in juvenile (t = 4.162, p < 0.001) but not in adult (t = 1.085, p = 0.289) animals; there is a developmental decrease for both avalanches starting in the BLA (t = 6.065, p < 0.001) and in the mPFC (t = 7.413, p < 0.001). Two-way repeated measures ANOVA, Holm-Sidak method for all pairwise multiple comparisons. (E) Normalized size distribution, calculated for local avalanches in the mPFC (*black*) and nLFP clusters in the mPFC that are followed by nLFP clusters in the BLA (*blue*) for juvenile (*left panel*) and young adult (*right panel*) males, averaged across animals. Shaded area corresponds to SEM. Grey rectangles and dashed vertical lines correspond to the significant differences (p < 0.05, Wilcoxon signed-rank test with FDR correction, q < 0.05). NLFP clusters that are part of PFC to BLA avalanches are characterized by a reduced likelihood of small avalanches and an increased likelihood of large avalanches compared to the local PFC avalanches. (F) Same as (E) for local avalanches in the BLA (*black*) and nLPF clusters in the BLA that are followed by nLFP clusters in the mPFC (*magenta*).

## Notes

### Competing Interest Statement

The authors have declared no competing interest.

